# Therapeutic potential of human mesenchymal stromal cell-derived mitochondria in a rat model of post-surgical digestive fistula: towards an energetic nano-biotherapy

**DOI:** 10.1101/2024.11.29.626014

**Authors:** Antoine Mariani, Augustin Guichard, Anna C. Sebbagh, André Cronemberger Andrade, Zahra Al Amir Dache, Christopher Ribes, Dmitry Ayollo, Mehdi Karoui, Gregory Lavieu, Florence Gazeau, Amanda K. A. Silva, Gabriel Rahmi, Sabah Mozafari

## Abstract

**Background:** Tissue regeneration heavily relies on cellular energy production, with mitochondria playing a crucial role. Dysfunctional mitochondria are implicated in various degenerative diseases, driving interest in targeting mitochondrial transplantation for tissue repair. Wound healing is highly compromised in gastrointestinal conditions resulting in fistula development, particularly after sleeve gastrectomy. Human mesenchymal stem/stromal cells (hMSCs) and their cell-free products such as mitochondria offer potential benefits due to their therapeutic properties on cellular energy production. Here we investigated the therapeutic advantage of hMSCs-derived mitochondria nano-biotherapy in a rat model of post-surgical fistula healing.

**Methods:** Viable and structurally intact mitochondria were isolated from hMSCs before exposure to human colonic epithelial cells (HCEC-1CT) culture or transplantation into a rat model of post-operative fistula.

**Results:** Our findings reveal significant dose-dependent improvement on cellular metabolic activity and ATP content of the recipient cells. Assessment of the external fistula orifice developed following post sleeve gastrectomy fistula, revealed a substantial healing in all transplanted rats compared to control group.

**Conclusion:** Our findings highlight the therapeutic potential of hMSCs-derived mitochondria in post-surgical fistula healing. This research contributes to advancing cell-free regenerative strategies for gastrointestinal conditions, offering new insights into mitochondrial-based therapies for enhancing wound healing and tissue repair.

## Background

Tissue regeneration is a highly dynamic and energy-consuming process associated with cellular energy production (1). ATP and various metabolites produced through energy metabolism serve as critical substrates for the biosynthetic pathways needed during tissue repair such as cell proliferation, differentiation, and extracellular matrix synthesis (2). Targeting energy production to enhance tissue repair has demonstrated promising results across various degenerative conditions (3). Mitochondria, known as the powerhouse of cells, play crucial roles in energy production and have been the focus of recent regenerative nano-biotherapies worldwide (4). While dysfunctional mitochondria lead to excessive reactive oxygen species production and cause oxidative damage in cells, functional mitochondria play important roles in the regulation of oxidative phosphorylation and generation of ATP or other metabolites. Mitochondrial function has been proposed as a key hallmark for intestinal epithelial cell plasticity, determining the regenerative capacity of intestine for wound healing (5, 6). The importance of mitochondrial function in gastrointestinal cell homeostasis has been widely highlighted by Rath and colleagues (7, 8).

Gastro-intestinal surgery as well as inflammatory bowel disease may lead to formation of digestive fistula, which is an abnormal structure connecting digestive organs or a digestive organ with skin. Fistula formation is among the most common complications after sleeve gastrectomy accounting for 1% to 5% of cases (9). With an increasing rate of obesity surgeries, this medical condition can be the cause of significant morbidity for patients, leading to sepsis or even death and also poses a significant managing challenge necessitating combined medical, radiological, endoscopic and sometimes surgical treatment modalities (10). Therefore, new perspectives including regenerative medicine are needed.

Different biotherapeutic strategies have been suggested to enhance tissue repair and regeneration in postoperative fistula including the use of stem cells or cell-free nano-biotherapies such as extracellular vesicles (EV) (11). Some regenerative effects of EVs are attributed to the presence or transfer of metabolically active products, enzymes, or entire mitochondria (12–15). Intercellular transfer of mitochondria has been proposed as a means of tissue revitalization (16). Physiologically, mitochondria can be transferred to the recipient cell via tunneling nanotubes, intercellular dendrites, vesicles cargos or via direct extrusion from donor cells and then internalization by the recipient cells (17). Moreover, it has been recently reported that blood contains between 200,000 and 3.7 million per milliliter of circulating cell-free respiratory competent/functional mitochondria in normal physiological state (18). Stephens et al., revealed that circulating mitochondria were able to enter rho-zero cells (with depleted mitochondrial DNA) and visualized using immunoelectron microscopic imaging (19). Inspired by these physiological mechanisms of intercellular mitochondria transfer, and recognizing that one of the primary mechanisms of EVs in tissue repair involves mitochondrial transfer, direct transplantation of healthy and functional mitochondria into injured tissues has emerged as a promising new therapeutic approach. It was first proposed by McCully et al. in 2009 via injection of healthy mitochondria into a rabbit model of ischemic heart (20). Moreover, the first clinical study was performed in 2017 in a pediatric population having extracorporeal membrane oxygenation (ECMO) secondarily to cardiac ischemia-reperfusion injury. They showed that 4 of 5 patients were successfully weaned off ECMO due to increased cardiac function after transplantation of autologous mitochondria into the myocardium (21). Other recently completed or ongoing clinical trials involving (human mesenchymal stromal/stem cell (MSCs)-derived) mitochondrial transplantation have been recently reviewed by Suh and Lee (22).

MSCs population is defined as progenitor cells capable of self-renewal, immunomodulation, differentiation anti-inflammatory, regenerative, pro-angiogenic, anti-apoptotic, anti-fibrotic potentials (23). These cells are available from different tissue sources and their properties enable them to treat various diseases. In the mechanisms underlying MSCs-based therapy, there is growing evidence showing the paracrine effects by release of microvesicles, exosomes or direct mitochondrial transfer into damaged cells (24, 25). MSCs based cell-free nano-biotherapies have recently attracted great attention mainly due to its broad advantages over cell therapy (26, 27). The strength of cell-free over cell biotherapies lies in the potential to mitigate the risks of tumorigenesis, unwanted cell differentiation, incorrect cell integration, vascular occlusion, and shelf-life gains, while retaining similar advantageous properties (28). Another key asset is the immune-privileged status of MSCs-derived products, which are devoid of major histocompatibility complex (MHC) suggesting their potential as “off-the-shelf” therapeutic modality allowing to consider allogeneic biotherapy (29). All these made MSC to be considered as an optimal cell source candidate for mitochondria cell-free biotherapy in different experimental conditions (12, 30).

Although the regenerative potential of mitochondria transplantation has been reported in different tissues, like in the brain (31), spinal cord (32), lung (13), heart (33, 34) or kidney (35), the beneficial effects of mitochondrial biotherapies for tissue repair specifically in the family of gastrointestinal diseases remain unexplored. The aim of this study is to investigate hMSCs-derived mitochondrial transplantation as a cell-free biotherapy for post-surgical fistula healing. We hypothesize that local administration of freshly intact mitochondria from human adipose tissue-derived MSCs would help regenerating damaged tissue in a post sleeve gastrectomy fistula model.

## Methods

### Human adipose tissue-derived mesenchymal stem cells (hMSCs) culture

Human adipose tissue-derived MSC (CellEasy batch number N°9297) were grown in T175 culture flasks using α-Minimal Essential Medium (α-MEM) supplemented with 10% fetal bovine serum (FBS) at 37 °C, 5% CO_2_ with 20 mL of media utilized to ensure complete flask coverage. The medium was refreshed every 3 days, and cells were grown until reaching 90% confluence before passaging.

To detach the cells, the culture medium was aspirated, and the flasks were washed with phosphate-buffered saline (PBS, Gibco, Thermofisher). The washed cells were then incubated with 0.05% trypsin ethylenediaminetetraacetic acid (EDTA, Gibco, Thermofisher) for 2 to 3 minutes at 37°C with 5% CO_2_. Mechanical detachment was facilitated by tapping the flasks. Following trypsinization, an equal volume of α-MEM with 10% FBS was added to neutralize the trypsin. The cell suspension was centrifuged at 300 g for 10 minutes, and the supernatant was discarded. Cell quantification and viability assessment were performed using the NucleoCounter® NC-200™ automated cell analyzer (NC-200, Chemometec). Only cells from passages 4 to 8 were utilized for mitochondria isolations.

### Isolation of mitochondria from hMSCs

A commercial mitochondrial isolation kit for cultured cells (ab110170, Abcam) (36–39) was used to isolate mitochondria from hMSCs. Initially, cells were pelleted via centrifugation at 300 g, followed by rapid freezing and thawing to weaken cell membranes. Subsequently, the cells were suspended in Reagent A at a concentration of 5.0 mg protein/ml (approximately 1 ml per 25 million cells) and incubated on ice for 10 minutes. Cell disruption was achieved using a sonication probe (Sonicator FB50, Fisher Scientific), with ultrasound applied in 3 cycles of 10-second bursts at 20% power, separated by 10-second intervals (total active phase of 30 seconds), while keeping the cell suspensions cooled on ice.

Next, the homogenate underwent centrifugation at 1,000 g for 10 minutes at 4°C to remove remaining cells, cell debris, and nuclei. The resulting supernatant (SN) #1, containing mitochondria, was collected, while the pellet was resuspended in Reagent B to the same volume as Reagent A. The cell rupturing step was repeated to release mitochondria from the remaining intact cells and the resulting supernatant SN #2 was collected, while the pellet was discarded. SNs #1 & #2 were combined, thoroughly mixed, and centrifuged at 12,000 g for 15 minutes at 4°C. The resulting pellet constituted the isolated mitochondrial mass was resuspended in ice-cold respiration buffer (RB): 250 mM sucrose, 5mM KH_2_PO_4_, 10 mM MgCl_2_, 20 mM HEPES (pH 7.2) and 1 mM EGTA (pH 8); quantified, dose adjusted, aliquoted and kept on ice until transplantation. For mitochondria characterization, the pellet was washed with Reagent C supplemented with Protease Inhibitors (11697498001, Roche), and aliquoted for the subsequent characterization, *in vitro* or *in vivo* experiments or stored at -80°C until the mitochondrial quality assays were conducted.

### Visualization of isolated mitochondria

MitoTracker Red CMXROS is a red-fluorescent dye that selectively stains intact mitochondria with its accumulation contingent upon mitochondrial membrane potential. Viability of freshly isolated mitochondria was assessed using MitoTracker Red CMXROS (M7512, Invitrogen) staining at 200 nM for 40 minutes followed by two washes with PBS 1X and immediate observation of mitochondria (resuspended in 10 µL in PBS, mounted onto slides) using fluorescence microscopy (EVOS M5000, ThermoFisher).

### Immunofluorescence imaging of isolated mitochondria

Immunofluorescence imaging of isolated mitochondria involved characterizing freshly isolated samples through mitochondria-specific immunostaining (following or not MitoTracker Red CMXROS staining). Initially, isolated mitochondria were incubated either with an Alexa Fluor® 488 Anti-TOMM20 (translocase of outer mitochondrial membrane 20 homolog) antibody [EPR15581-39] - Mitochondrial Marker (ab205486, abcam) or with a primary rabbit anti-mouse anti-TOM20 antibody (11802-1-AP, Proteintech) for 30 minutes followed by 30 minutes incubation in the dark with a secondary antibody (Goat anti-Rabbit IgG) conjugated with green fluorescent AlexaFluor 488 (A-11008, ThermoFisher). TOM20 antibody specifically binds to the TOM20 complex located in the mitochondrial outer membrane. Samples were washed twice with PBS 1X and underwent centrifugation (12000 g, 15 minutes, 4°C) at each washing step. Visualization and imaging were achieved using a Cell Insight CX7 LZR automated confocal spinning-disk fluorescence microscope (ThermoFisher Scientific) or EVOS M5000 microscope (ThermoFisher).

### Western blot analysis of mitochondria protein markers

Mitochondria obtained from hMSCs underwent homogenization and lysis in ice-cold 30 mM Tris-EDTA buffer, pH 7.2, supplemented with 1 mM DTT, 1% (v/v) Triton X-100, 10% (w/v) anti-phosphatase cocktail, and 14% (w/v) anti-protease cocktail (11697498001, Roche), for 30 minutes on ice. Homogenate was gently mixed on an agitating plate for 2 minutes at ice-cold temperatures. The proteins were quantified using a Bradford assay. From each sample 20 µg of total protein was loaded and separated on 4–20% Mini-PROTEAN® TGX™ Precast Protein Gels (Bio-Rad), followed by electroblotting onto membranes of Trans-Blot Turbo Midi 0.2 µm PVDF Transfer Packs (Bio-Rad). Subsequently, the membranes were rinsed in TBS-Tween 20 buffer (TBST) and blocked for 1 hour in Every-Blot Blocking Buffer (BioRad) before being washed with TBST.

For antibody incubation, the membranes were then incubated overnight at 4°C with either: (i) rabbit anti-TOM20 (1:5000, 11802-1-AP, ThermoFisher); (ii) total oxidative phosphorylation (OXPHOS) Human WB Antibody Cocktail (MS601) (1:700, ab110411, Abcam); (iii) Membrane Integrity WB Antibody Cocktail (MS620) (1:250, ab110414, Abcam); or (iv) rabbit anti-GAPDH antibody (1:2500, ab9485, Abcam) as a loading control. Following washing, the blots were incubated with the corresponding goat anti-rabbit horse-radish peroxidase (HRP)-linked Antibody (1:2000, 7074, Cell Signaling) or goat anti-mouse HRP-conjugated secondary antibody (1:2000, 7076, Cell Signaling) for 2 hours at room temperature.

Peroxidase activity was revealed using a chemiluminescent detection kit (ECL Plus substrate, GE Healthcare). Protein expression visualization and acquisitions were performed using the Syngene PXi (UK), with documentation accomplished using GeneSys image capture software (UK). Post-spin supernatant was loaded as a control at 20μg for all the experiments.

### Interferometric light microscopy (ILM) measurements of isolated mitochondria using VideoDrop

Measurements of isolated mitochondria using ILM with Video-Drop technology were conducted to determine their concentration and average hydrodynamic radius. Drops of 7 µL each were dispensed onto a round cover slip using a micropipette, followed by positioning the stage towards the objective. The QVIR 2.6.0 software (Myriade, Paris, France) was employed to illuminate the sample using a 2 W blue LED light.

The concentration of particles was determined by the number of detected particles within the measured volume, which is influenced by both microscope parameters and particle size. The hydrodynamic radius (size) was estimated by tracking the motion of imaged particles in recorded movies, as they undergo thermal agitation. All detected particles contribute to the number density of particles per milliliter (concentration).

Tracking was initiated with a minimum of 300 particles, with up to 50 videos analyzed, each containing 100 frames. A minimum saturation of 92% was considered, and a relative thresh-old of 4.2, as specified by the manufacturer, was applied for detection. Macroparticle detection was enabled by setting a minimum radius of 10 and a minimum number of hot pixels of 150, along with drift compensation. To ensure accuracy, signals from PBS 1X were monitored and consistently found to be below the detection limit.

### Electron Microscopy

#### Scanning Electron Microscopy (SEM)

Samples were mounted on aluminum stubs (32 mm diameter) with carbon adhesive discs (Agar Scientific, Oxford Instruments SAS, Gomez-la-Ville, France) and visualized by field emission gun scanning electron microscopy (SEM FEG) as secondary-electron images (2 keV, spot size 30) under high-vacuum conditions with a Hitachi SU5000 instrument (Milexia, Saint-Aubin, France). SEM analyses were performed at the Microscopy and Imaging Platform MIMA2 (INRAE, Jouy-en-Josas, France) DOI: MIMA2, INRAE, 2018. Microscopy and Imaging Facility for Microbes, Animals and Foods.

#### Transmission electron microscopy (TEM) - Negative-staining

Isolated mitochondria (5 µL) were applied directly onto a carbon film membrane on a 300-mesh copper grid, followed by staining with 1% uranyl acetate solution dissolved in distilled water. The grids were subsequently air-dried at room temperature. Examination of the grids was performed using a Hitachi HT7700 electron microscope operating at 80 kV (Elex-ience), and images were captured with a charge-coupled device camera (AMT).

#### Transmission electron microscopy (TEM) - Cross-sectioned method

Mitochondrial suspensions (45 μL) were subjected to centrifugation at 12,000 ×g for 15 minutes, after which the supernatant was carefully removed, and the resulting pellets were fixed with 2% glutaraldehyde in 0.1 M sodium cacodylate buffer at pH 7.2 for 1 hour at room temperature. Following removal of the supernatant, the pellets underwent three washes in PBS 1X. Subsequently, samples were contrasted with 0.2% Oolong Tea Extract (OTE) in cacodylate buffer, postfixed with 1% osmium tetroxide containing 1.5% potassium cyanoferrate for 1 hour. After an additional three washes in PBS 1X, the pellet was dehydrated through a graded series (30% to 100%) and gradually substituted in a mixture of ethanol-epon, finally being embedded in 100% Epon (Delta microscopie, France) over a 24-hour period. The pellet underwent a final embedding in Epon, cured at 37°C overnight, and further hardened at 65°C for two additional days.

Ultrathin sections (80 nm) were then collected onto 200-mesh copper grids, stained with 2% uranyl acetate in 50% methanol for 10 minutes, and counterstained with 1% lead citrate for 7 minutes. The grids were examined using a Hitachi HT7700 electron microscope operated at 80 kV (Milexia – France), and images were acquired with a charge-coupled device camera (AMT). This analysis was conducted at MIMA2 MET – GABI, INRA, Agroparistech, 78352 Jouy-en-Josas, France.

### *In vitro* evaluation of the effects of mitochondria on the metabolic activity of starved HCEC-1CT cells

Cells of the HCEC-1CT colon epithelial progenitor cell line were obtained from Evercyte. This cell line was immortalized from non-tumoral adult human colon biopsies. HCEC-1CT were routinely cultivated at 37°C with 5% CO_2_ in ColoUP medium, composed of four parts DMEM (Gibco, 31966-021) and one part M199 (Gibco, 31150-022), 2% Cosmic Calf Serum (Hyclone, SH30087), 20 ng/ml hEGF (Sigma, E9644), 10 µg/ml Insulin (Sigma, I9278), 2 µg/ml Apotransferrin (Sigma, T2036), 5 nM Sodium Selenite (Sigma, 214485), 1 µg/ml Hydrocortisone (Sigma, H0396). Cells from passages 10 to 58 were utilized for the subsequent experiments.

HCEC-1CT cells were seeded in 96 well plates (VWR 734-2327) at 5000 cells per well, then amplified for 48h in ColoUp medium. Differentiation was induced by 120h culture in a differentiation medium composed of four parts DMEM and one part M199, 0.1% Cosmic Calf Serum, 1.25ng/ml hEGF, 10µg/ml Insulin, 2µg/ml Apotransferrin, 5nM Sodium Selenite, 1 µg/ml Hydrocortisone, 5µM 6-bromoindirubicin-3-oxime / GSK-3 inhibitor (Sigma, 361550-1MG) and 100 µg/ml Primocin (InVivogen, ant-pm-05).

After differentiation, the cells were washed and starvation was induced by shifting to basal medium composed of four parts DMEM and one part M199 with no further supplementation. Three independent experiments were performed to isolate fresh-mitochondria from hMSCs. HCEC-1CT cells were then treated for 48h with either freshly-isolated hMSC mitochondria suspended in RB (at different doses of 10^8^-3.33×10^10^ particles/mL in the well) or an equivalent volume of RB (negative control). The positive control group was cultivated in fresh differentiation medium instead of basal medium, and treated with RB. Metabolic activity of the starved HCEC-1CT cells treated or not with freshly-isolated mitochondria (three independent batches) was evaluated at 24h and 48h through the Alamar Blue Assay (Invitrogen, DAL1100). To analyze the metabolic activity, in brief, cells were first washed in PBS, then incubated for 3h in Alamar blue reagent diluted 10 folds in white DMEM (Gibco, 31053-028). Metabolic activity was then estimated by the rate of reduction of resazurin into resorufin in viable cells as evaluated through fluorescence measurements (540/590nm, integration time 400ms, SpectraMax® iD3 Microplate Reader, Molecular Devices).

Mitochondria staining was performed to visualize mitochondria content of the HCEC-1CT cells 48h after *in vitro* treatment. Briefly, the cells were washed and stained with Hoechst at 200nM and Bio Tracker™ 633 Red Mitochondria Dye at 100nM (Sigma-Aldrich SCT137) for 30min at 37°C. The cells were then washed and kept in PBS. Sixteen fields per well and one well per condition were imaged with a Cell Insight High-Content Screening CX7 LZR automated confocal spinning-disk fluorescence microscope (Thermo-Fisher Scientific) at 20X magnification before analysis.

To further verify the cell metabolic activity following exposure to isolated mitochondria, the ATP content of a subset of wells was measured using the ATP Bioluminescence Assay Kit (Abcam ab113849) according to the supplier’s instructions. We then evaluated whether the cells’ ATP contents at 48h correlated with their metabolic activity measured by the Alamar blue assay through computation of Pearson’s correlation coefficient after checking whether the assumption of normality could reasonably be made then linear regression analysis, using the GraphPad Prism software (version 8.0.2 and 10.0.2).

### Rat model of gastro-cutaneous fistula

All experiments were approved by the animal care and use committee in France and the Ministry of Higher Education and Research (APAFIS #35921-2022010121242601). A total of 20 female 11-week-old Wistar rats were divided into 2 groups: a control group (n = 10), an experimental group (n = 10) with a mean weight of 221 g (range 212g-235g). Minimization method was used to ensure groups were comparable in terms of gender, weight and age. The sample size was determined based on our previous studies evaluating the therapeutic potential of hMSCs-derived EVs or hMSCs in post operative fistula healing model (40).

The animals went through a 7-day acclimatization period with water and food ad libitum. They were housed in the laboratory animal room, in cages, with regulated temperature, ventilation, and respecting light–dark cycles. Anesthesia was performed under 2% of isoflurane and analgesia was induced with 0.05mg/kg of buprenorphine. After a midline laparotomy of 2-3 cm, the entire greater curve of the stomach was identified. Pylorus and esogastric junction were identified and gastrosplenic ligament sectioned. The “sleeve” gastrectomy was performed using a linear stapler, from 5mm of the pylorus to the distal part of the rumen, carefully avoiding stenosis of the remnant gastric tube. The upper part of the rumen was passed with a staggered opening through the abdominal muscle and the subcutaneous space, forming a 5 mm-long tract. Four stitches (PDS 4/0) were used to attach the stomach to the skin at the site of the incision on the left flank of the rat, creating a 5mm large gastro-cutaneous fistula model. Postoperative analgesia was performed.

### Mitochondria transplantation

Fourteen days after surgery, animals (n = 10) were anesthetized using 2% isoflurane, and were locally injected at the lesion site with 0.6 mL of freshly isolated mitochondria (9.9 x 10^10^ particle/mL per rat, equal to 800µg mitochondria protein, generated by 17×10^6^ hMSCs) resuspended in mitochondria respiration buffer. Each animal received 4 local injections of 100 µL each of fresh mitochondria (around the external orifice of fistula plus 200 µL injected directly inside the fistula tract wall). Mitochondria from three independent isolation were used. Control group similarly received 0.6 mL of PBS or respiration buffer (n = 10). With one animal in the control group died following the surgery, 19 animals (in total) were finally followed-up until the end of the experiments.

### Preclinical follow-up

Rats (n=10 for Mito vs n=9 for control groups) were daily followed up to 45 days post operation (DPO) and clinically assessed at DPO 0, DPO 14, DPO 21, DPO 28 and DPO 45 corresponding to 30 days after treatment). Animals were anesthetized every week, using 2% isoflurane, for fistula measures using a graduated ruler, and pictures were taken for external control. The presence of feces at the fistula orifice, indicative of positive fistula output, was evaluated macroscopically. The absence of feces indicated negative fistula output.

### Histological analyses

At D60, rats were sacrificed, under 2% isoflurane anesthesia, using intracardial injection of thiopental. The fistula site as well as its periphery (1 cm length, 2 specimens each per rat) were collected and transferred to a formalin solution. Specimens were embedded in paraffin and sectioned perpendicular to the center of the fistula to obtain thin tissue sections of 7 μm, which were stained with hematoxylin and eosin (HE) and Sirius Red (fibrosis assessment). Slices were analyzed with an optical microscope (Leica DMIL). Histological analysis was performed by an external pathologist (CB), blinded to treatment allocation. All slices were also digitally scanned (Digitiser Hamamatsu Photonics®, Massy, France) and analyzed with dedicated software (Halo) for fibrosis quantitative analysis.

### Statistics

The results were presented as means ± standard deviation for continuous variables, and as percentages for categorical variables. Fischer’s exact test was carried out for comparisons between categorical variables and the nonparametric Mann–Whitney test was used for non-paired continuous variables. An estimation of the p value by the Chi-square test was carried out for the comparison concerning the number of cases per group. Pearson’s correlation coefficient was calculated, and linear regression analysis was conducted for the *in vitro* assessment of ATP content and Alamar Blue metabolic activity. A two-way ANOVA, followed by Sidak’s multiple comparisons test, was employed to evaluate differences in the fistula orifice diameter over time between the treated and control groups. Statistical analysis was conducted using SAS version 9.1 (SAS Institute, Cary, NC, USA) or GraphPad Prism v.8.0.2 software (Graphpad Software, La Jolla, CA, USA). The notations for the different levels of significance are indicated as follows: *p < 0.05, **p < 0.01, ***p < 0.001, ****p < 0.0001. For all the experiments, a p-value inferior to 0.05 is considered significant.

## Results

### Viable and structurally intact mitochondria are isolated from hMSCs

The mitochondria from hMSCs were fully characterized through various assays. Immunofluorescence staining revealed robust expression of TOM20, a mitochondrial outer membrane marker (Figure 1A). Further assessments through co-labeling with MitoTracker Red as well as TOM20 (Figures 1Bi-iii) demonstrated the presence of viable mitochondria. Scanning electron microscopy (SEM) imaging showed multiple rounds to oval-shaped structures ranging from 0.2 to 2 µm in size with intact outer surfaces (Figure 1C), corroborated by TEM analyses of negative staining or cross-sectioned methods depicting the presence of multiple mitochondria with inner membranes and a diverse range of diameters spanning from 0.1 to 1.2 µm (Figures 1Di-Dii).

**Figure 1.**
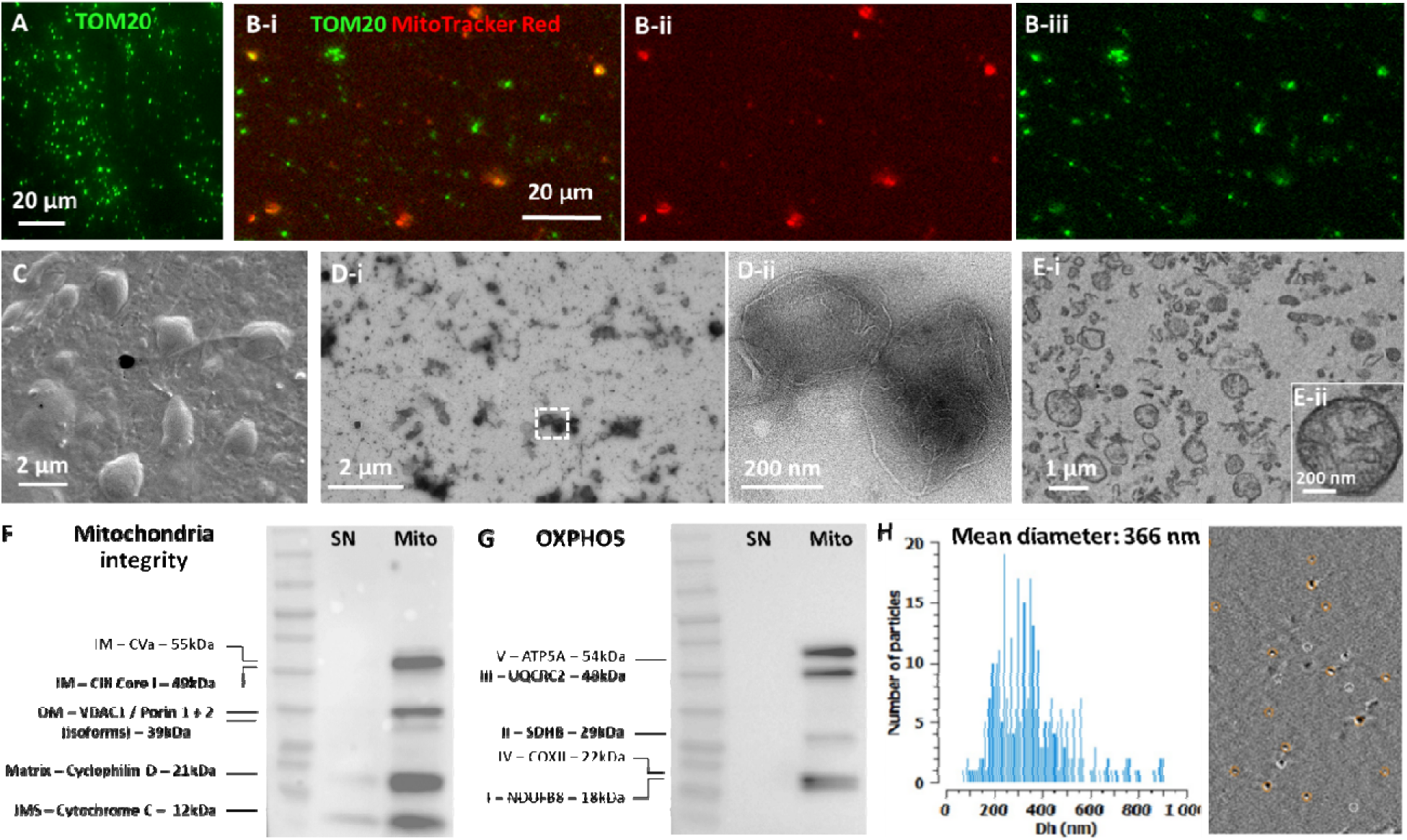
Characterization of human MSCs-derived mitochondria. (A) TOM20 (green) staining and (Bi-iii) MitoTracker red/TOM20 colabling of freshly isolated mitochondria indicating isolation of numerous viable mitochondria from hMSCs. (C-D) Scanned electron microscopy (SEM) and transmission electron microscopy using negative staining (Di-Dii) as well as cross-sectioned methods (Ei-Eii) depicting structurally intact mitochondria of various diameter ranging from 100-1200 nm. (F) Quality control analysis of isolated mitochondria by Western blotting of mitochondria membrane integrity markers and (H) human OXPHOS complexes (I-V) showing co-expression of mitochondrial structural and functional proteins in mitochondria preparations (Mito) compared to their corresponding supernatant (SN) loaded with the same amount of protein (Uncropped western blots can be found in the Supplementary Figure 1). (E) ILM of mitochondria samples using Videodrop particle counter showing the size distribution graph with a mean size of 366 nm and a representative interferometry image (depicting the particles in white circles appeared for a moment or the tracked particles shown in circles with orange color). IM: inner membrane; OM: outer membrane; IMS: intermembrane space.

Quality control assessment of the isolated mitochondria was conducted through a series of Western blot analyses. For each preparation, the corresponding supernatant (SN) was used as a control to facilitate sample characterization. An equal amount of 20 µg protein was loaded for both the mitochondria and SN samples (Figure 1F). The co-presence of five key mitochondrial membrane integrity markers (Figure 1F), as well as five markers of the human oxidative phosphorylation (OXPHOS) complexes I-V in mitochondria preparations (a total of 10 markers), was confirmed by Western blot analysis for each independent mitochondria isolation experiment.

Mitochondrial size distribution analysis was performed using interferometric light microscopy (ILM) by videodrop particle counter. The mean size of isolated mitochondria was determined to be 366 nm, consistent with previous literature and TEM analysis (Figure 1E).

### hMSC mitochondria promote the metabolic activity of starved colon cells *in vitro*

Next, we evaluated the effect of freshly isolated hMSC-derived mitochondria on the metabolic activity of starved HCEC-1CT colon cells assessing their metablic activety via Alamar blue assay, ATP concentration via ATP assay as well as mitochondria content using mitochondria memberane potential indicator, Biotracker red, staining and quantification. Starvation was induced through cultivation in basal medium and the cells were treated with mitochondria suspended in RB. RB alone was used as a negative control. Non-starved cells treated with RB were used as a positive control.

The metabolic activity of the cells, assessed through the Alamar blue test was measured at 24h and 48h. While no metabolic activity could be detected for the mitochondria alone, Alamar or white DMEM, when normalized to positive control (non-starved cells treated with RB), a remarkable dose-response effect was observed following treatment with mitochondria of different concentrations of respectively 3.35E10 (p value < 0.01) and 7.07E9 (p value < 0.001) at 24h (Figure 2A) or 3.35E10 (p value < 0.001), 7.07E9 (p value < 0.001) and 2.83E9 (p value < 0.01) at 48h compared to negative control (Figure 2B).

**Figure 2:**
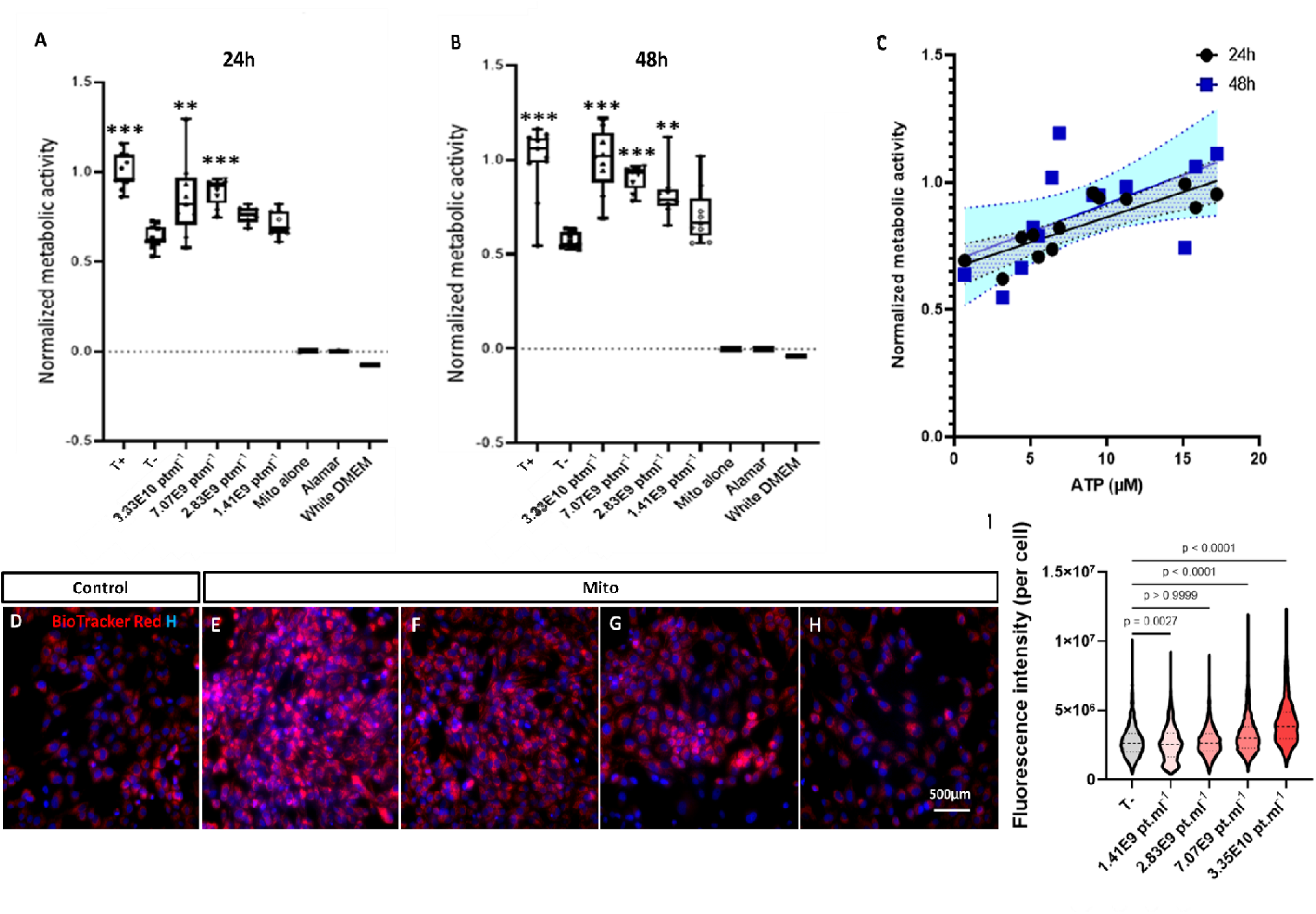
Metabolic activity of HCEC-1CT colonic cells following in vitro mitochondria transplantation. (A and B) Alamar results of the metabolic activity of cells exposed to fresh mitochondria at different doses compare to negative control (non-starved cells treated with RB) at 24h (A) and 48h (B) normalized to positive control. (C) Normalized metabolic activity evaluated through the Alamar blue test as a function of the ATP concentration – Linear regression analysis. The ATP contents of HCEC-1CT cells was evaluated at 48h in a representative subset of 13 wells. Data are shown as individual values with the best-fit curve and its 95% confidence bands (shaded area). (D to H) Fluorescence images of stained cells (Red: Biotracker 663; Blue: Hoechst) 48h after treatment with different doses of mitochondria; (D) Negative control; (E) Mitochondria condition 3.35E10 pt.ml^-1^; (F) Mitochondria condition 7.07E9 pt.ml^-1^; (G) Mitochondria condition 2.83E9 pt.ml^-1^; (H) Mitochondria condition 1.41E9 pt.ml^-1^. (I) Repartition of Biotracker 663’s fluoresce intensity per cell for each condition; p values were calculated using Kruskal-Wallis non-parametric test. ****: p<0.0001, ***: p<0.001, **: p<0.01, *: p<0.05); T-: negative control; T+: positive control.

Interestingly, Alamar results both at 24h and 48h significantly correlated with HCEC-1CT ATP contents in a representative subset of 13 wells at 48h. Metabolic activity as a function of the ATP contents at 24h and 48h are shown in Figure 2C. The Pearson coefficient was of 0.837 with a 95% confidence interval of [0.529; 0.950] at 24h, and 0.581 with a 95% confidence interval of [0.043; 0.858] at 48h. Linear regression analysis revealed significant deviation from 0 and non-significant deviation from linearity at both time-points, with a R square of 0.6997 at 24h and of 0.3370 at 48h.

To assess the mitochondria content of each cell condition after treatment with freshly isolated mitochondria, cells were counterstained with Hoechst and BioTracker 663 red at 48 hours. BioTracker 663 red is an indicator of mitochondrial membrane potential commonly utilized in live cell imaging to detect cell viability or metabolic activity (Figure 2D-H). While the same number of cells were plated to evaluate the metabolic activity of cells exposed to mitochondria with different doses, the cells survival (detected by higher cell density) seemed to be clearly and does-dependently higher for the mitochondria-treated vs control conditions (Figure 2D-H). The fluorescence intensity of BioTracker 663 Red, associated to viable mitochondria per cell was measured for wells treated with different mitochondria doses and compared to that of negative control. Results revealed dose-dependent intensities of 4.06×10^6^ ± 3.33×10^4^ (p value < 0.0001) and 3.21×10^6^ ± 2.46×10^4^ (p value < 0.0001) at doses of respectively 3.35E10 and 7.07E9 (Figure 2I). The conspicuous dose-response effect observed with the higher fluorescent intensity for the condition treated with the higher mitochondria dose (and vice versa) is visualized in Figure 2D-I.

### Post-surgical gastro-cutaneous fistula model in rats

A surgical gastro-cutaneous following “sleeve” gastrectomy was performed by gastrostomy at day 0 (experimental design is depicted in Figure 3). Following the procedure, all fistulas featured an external orifice diameter of larger than 4mm. There was no complication related to surgery. The mean of weight before surgery was 228.4 +/- 14g. Evolution of weight during follow-up showed a mean range started from 218.3 +/- 18g at day 14, right before treatment, to 232.7 +/- 28g at the end of the follow-up, just before sacrifice. No difference in animal weight was observed between the control and treated groups. All fistulas were permeable (persistent opening) 14 days after the initial procedure with a mean diameter of 4.1 +/- 1.3mm, indicating the absence of spontaneous closure before treatment.

**Figure 3:**
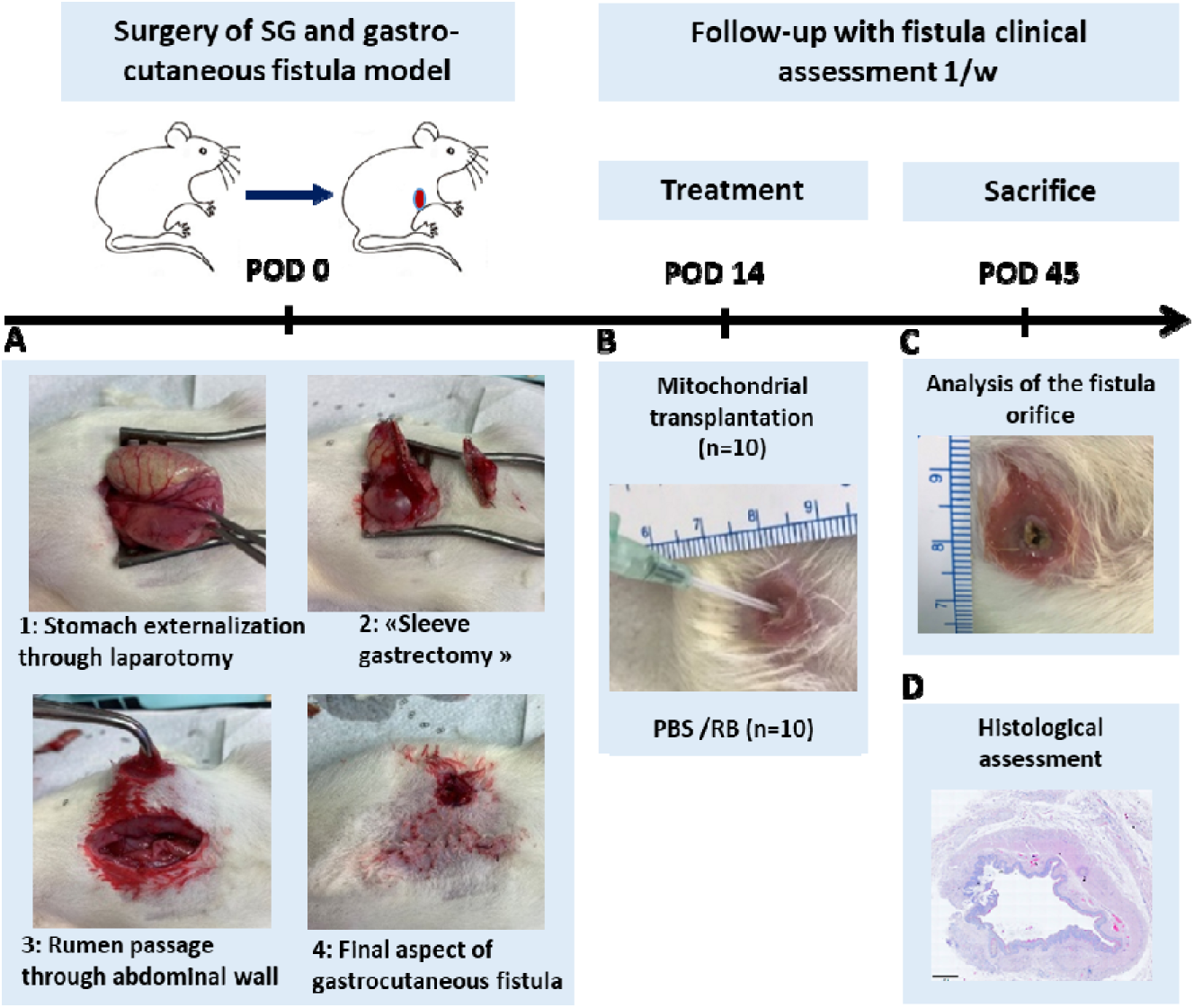
Design of the in vivo experiments. Study timelines and sample size are displayed. (A) Surgical procedure for the gastro-cutaneous fistula model induction from laparotomy (step 1), “sleeve gastrectomy” (step 2), rumen passage through abdominal wall (step 3) to final gastrocutaneous fistula formation (step 4) resulting in stomach communication to the skin in rats. (B) Percutaneous and intraluminal administration of the mitochondria via the fistula external orifice at days 14 post surgery. (B and C) Weekly follow-up with clinical assessment of animals and analysis of the fistula orifice diameter over time (at POD 0, POD 14, POD 21, POD 28 and POD 45). (D) Histological assessment of the postmortem tissues at POD 45. POD: post-operation day.

### Mitochondrial transplantation

Freshly collected mitochondria suspended in respiration buffer were grafted between 35 and 55 minutes after sampling in the mitochondrial group at day 14 post operation (14 POD). Control group similarly received PBS (n=5) or respiration buffer (n=5) with no significant difference between them during the entire experiment (at all 5 different clinical evaluation). Animals were clinically evaluated every week following transplantation until the end of study. No evident signs of allergy or toxicity were observed during the entire follow-up. At 21 DPO (7 days post transplantation), although no fistula closure was observed in the animals (Figure 4A-B), the mean diameter of external orifice was significantly reduced to 1.7 ± 1.09 mm in the mitochondrial group (n = 9 animals) versus 4.1 ± 1.19 mm in the control group (n = 10 animals, p value <0.0003). At D45, complete closure of the fistula occurred in 2 cases (22.2% of animals) with the mean orifice diameter of 0.88 ± 0.6 mm (n = 9 animals) in the mitochondrial group compared to 4.3 ± 1.82 mm (n = 10 animals, p<0.0001) in the control group with 0% complete orifice closure (Figure 4B). The frequency of no fistula output (featuring the absence of feces, Figure 4C) was significantly (p= 0.0054, Fisher test) different when comparing the control group with 20% (2/10 animals) vs mitochondria group with 88.8% (8/9 animals). Moreover, 8 of 9 cases (88.8%) showed an orifice diameter ≤1mm in the mitochondrial group versus only 1 of 10 (10%) in the control group. Histological analysis of the post mortem tissues was carried out at 45 POD to assess inflammation and fibrosis (Figure 4D-F). An extended inflammatory zone surrounding the external orifice was found in the control group in comparison with the mitochondrial group. Qualitative analysis showed the presence of a persistent ulceration (Figure 4D) in 100% of the control samples (10/10 animals), which was significantly (p= 0.0001190, Fisher test) higher than that in the mitochondrial group (observed only in 1/9 animals, 11 %). In the fibrosis quantitative analysis, we found that intensity of fibrosis was not different between groups (with 39% (+/-7) of fibrosis intensity in the slices of the control group vs. 42% (+/-7.5) in the experimental group) but the corresponding area was nearly two times higher in the control group (with 34.3 mm^2^ of fibrosis analysis area in the control group vs. 18.3 mm^2^ in the experimental group, p=0.03, Figure 4E).

**Figure 4:**
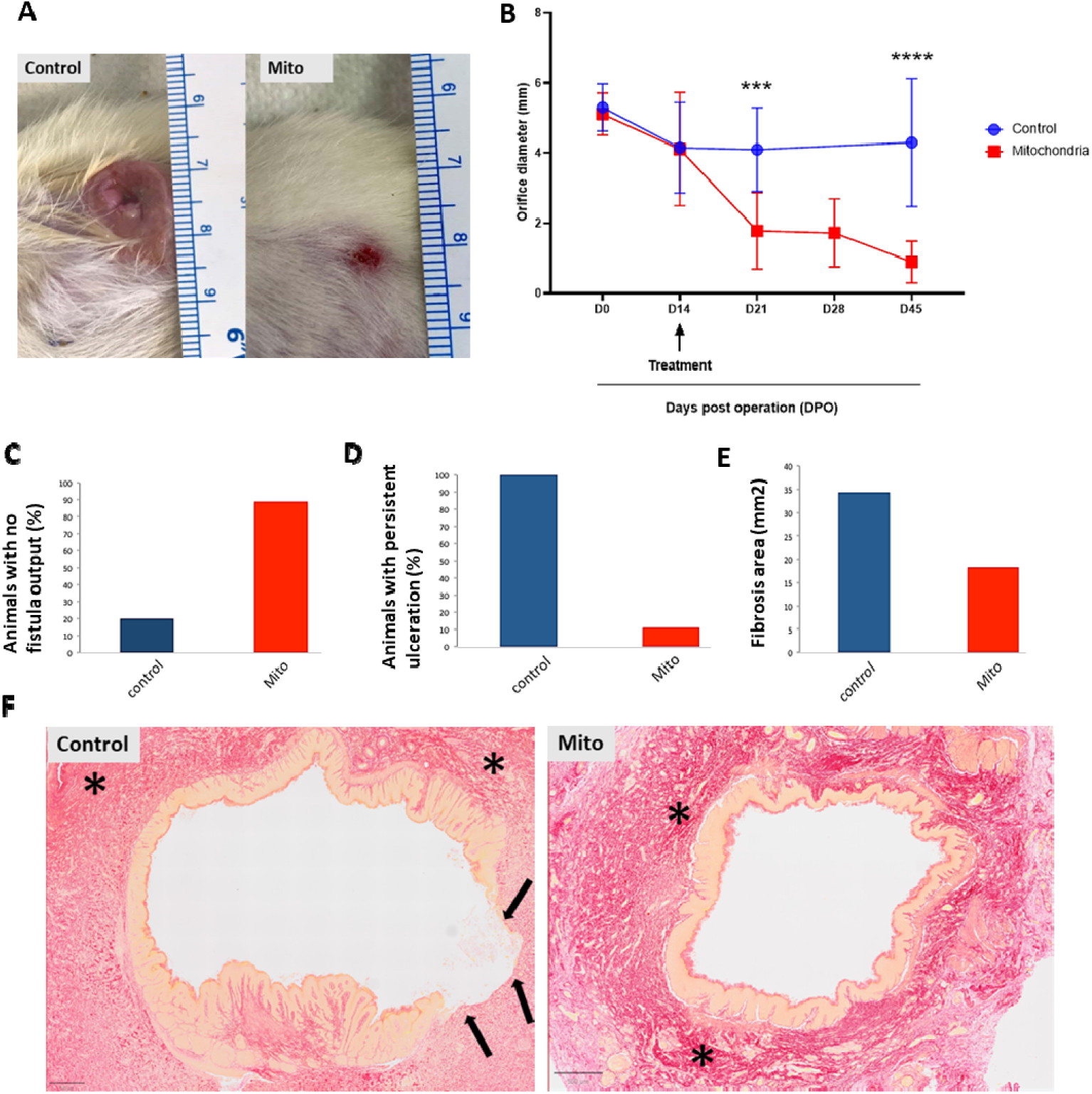
Potency evaluation in vivo following mitochondria transplantation. (A) Clinical assessment of the fistula orifice diameter performed on a weekly basis at D14, D21, D28 and morphological correspondence D45 post operation. (B) The orifice diameter is significantly reduced over time following local transplantation of hMSC-derived mitochondria (n=9) compared to the control group (n=10). The major reduction in orifice size is observed at D21 corresponding to 7 days after treatment. (C) Preclinical evaluation at day 45 indicating the percentage of animals per group featuring the absence of feces (no output) at the external fistula orifice as well as percentage of animals per group featuring persistent ulceration (D). Histology analysis (Sirius red staining) showed a decreased fibrosis area in the mitochondrial group (E) with corresponding slices (F). The main fibrosis regions in red staining were delimitated by *. The epithelium was identified by an orange staining. Important epithelial damage was observed for the control group and its extent was indicated by black arrows. Two-way ANOVA, Sidak’s multiple comparisons test was used for the statistical analysis of the differences. *** = p=0.0003, **** = p<0.0001, DOP: days post operation.

## Discussion

In the current era of obesity pandemic, laparoscopic sleeve gastrectomy (LSG) is one of the most frequently performed bariatric surgery worldwide (41). It is a relatively safe procedure with a postoperative fistula rate varying between 1% and 5% of cases (9). This complication could be particularly difficult to manage necessitating combined medical, radiological, endoscopic and sometimes surgical treatment modalities. Therefore, new perspectives including regenerative medicine are needed. In this regard, different cell-based or cell-free based biotherapies have been investigated. Due to the vital role of energy metabolism in wound healing and the importance of restoring mitochondrial biogenesis in gastrointestinal cell plasticity and homeostasis, especially in treating ulcerative colitis and Crohn’s disease, here in we investigated a mitochondria-based cell-free biotherapy for treatment of the post-surgical fistulas.

Cell therapeutic approaches have already been employed in rat models of gastric leakage. Benois et al. recently tested local administration of MSCs and platelet-rich plasma (PRP) and showed improvement of leak closure after this therapy (42). A first-in-human series with autologous MSCs-PRP endoscopic administration in 12 patients having gastric staple line fistula after LSG showed encouraging gastric leak healing results as well (43). Moreover, the biotherapeutic potential of cell-free MSC-derived EVs administered locally for the management of post-surgical fistula model of esophago-cutaneous fistula in pigs or post-surgical colo-cutaneous fistula in rats have been demonstrated recently by our team (40). Interestingly, analysis of MSCs-EVs revealed enrichment in proteins related to energy production and the presence of mitochondria (13, 44). Of note, MSC-derived EVs rescued mitochondrial dysfunction and improved clinical scores via extracellular vesicle mitochondrial transfer, in models of acute respiratory distress syndrome (ARDS) (13) or clinically relevant lung injury models (45). Mitochondria-rich extracellular vesicles from autologous stromal cell-derived cardiomyocytes restored energetics of ischemic myocardium (46) and were able to rescue patient-specific cardiomyocytes from toxic injury (15).

The aim of mitochondria-based therapies is to replace intracellular abnormal mitochondria with healthy mitochondria isolated from normal cells, resulting in the repair of damaged tissue and recovery of cell function (47). Inspired by the physiological means of intercellular mitochondria transfer, application of intact mitochondria isolated from healthy cells (referred to as mitochondria transplantation) has recently become an attractive nano-biotherapy strategy in regenerative medicine.

Different tissue sources for mitochondrial transplantation have been considered in different studies (48). Preble et al. employed a quick isolation protocol via differential filtration and obtained autologous mitochondria from muscles with the mean diameter of 380 nm measured by Coulter counter (49). Another source of healthy mitochondria includes platelets, which recently have attracted attention because they are abundant in blood that they can be simply acquired by venipuncture (50, 51). In addition, the isolation of mitochondria from platelets is relatively less invasive than that from muscles. Here, we focused on isolation of allogenic mitochondria from human MSCs. This choice was assumed from our experience of MSC-derived EVs and the potential development of an “off the shelf” solution due to the advantageous characteristics linked to the hMSCs (52, 53). More specifically, data suggest that MSCs could offer a promising therapeutic option for gastrointestinal conditions (54, 55). Our results show that structurally intact and viable mitochondria with conserved membrane potential and of high quality co-expressing 10 structural and functional markers - associated with mitochondria membrane integrity and oxidative phosphorylation complexes - are isolated from hMSCs. The mean diameter of hMSCs-derived mitochondria measured with ILM was 366 nm which is in agreement with that in the previous publications on muscle or adipose tissue derived hMSCs-mitochondria (56).

Next, to directly evaluate the metabolic effect of naked mitochondria on the gastrointestinal cells, freshly isolated hMSCs-derived mitochondria were introduced to a simplified *in vitro* model of HCEC-1CT cells under starvation. Starvation is known to modify the cellular metabolic activity *in vitro* (57). We evaluated the dynamic metabolic activity of starved HCEC-1CT cells 24h and 48h following hMSCs-derived mitochondria transplantation. Mitochondria treatment induced a significant dose-dependent effect on the metabolic activity of cells in 100% of *in vitro* experiments. This can be linked to the internalization of mitochondria by HCEC-1CT cells or the trophic effect of the co-isolated proteins. Internalization of the transplanted mitochondria by the recipient cells has been frequently reported before using different mitochondria membrane probes of different florescence wavelengths (48, 58). To address if the observed metabolic effect is linked to the energy production, the intercellular ATP content of the HCEC-1CT cells was measured in parallel at similar time points. Our data revealed a significant correlation between the dynamic metabolic activity measured by Alamar test and ATP concentration, that is, the more concentrated mitochondria transplanted the higher is the metabolic activity linked to the energy production revealed by an increase in the levels of cellular ATP. Similar results were reported by others using different recipient cells (59).

Furthermore, when the cells were stained with mitochondria membrane marker Biotracker red, a similar dose-dependent effect was observed. Although this experiment cannot confirm the internalization of exogenous mitochondria by the recipient cells, a clear difference in the total number of mitochondria per cell was observed as a function of dose.

We also showed that direct injection of allogenic mitochondria in the rat model of post-operative fistula was very well tolerated up to 31 days after delivery (45 DPO) without any allergic effect. Our clinical and histological data show a significant decrease in the fistula orifice diameter following hMSCs-derived mitochondria transplantation in 100% of grafted animals with the maximum effect observed at 7 days post transplantation (21 DPO) with no significant difference in orifice diameter between 21, 28 or 45 DPO. Based on this observation and the results of our previous studies using EVs in the same animal model (40), we have considered 45 days as the final time point for this study. Several *in vivo* studies have demonstrated rapid and sustained therapeutic effects of exogenous mitochondria in various animal models. For example, McCully et al. (2023) reported that transplanted mitochondria were detectable in myocardial cells, with significant therapeutic effects observed at 2, 4, 8, and 24 hours, as well as at 28 days post-delivery, with no adverse effects noted across different doses in a heart ischemia-reperfusion model (48). Additionally, another recent study showed that exogenous mitochondrial transplantation improved survival and neurological outcomes 72 hours after resuscitation from cardiac arrest (60). These findings suggest that mitochondrial grafts can exhibit efficacy shortly after transplantation. Nevertheless, future studies should investigate different doses of hMSC-derived mitochondria transplanted at various time points post-surgery to determine the optimal therapeutic window and assess any potential long-term adverse effects in digestive fistula model(s). Concerning the therapeutic potency, experimental results of mitochondrial transplantation have shown a reduction of the infarct size in cardiac ischemia-reperfusion injury, in rat and swine models (61). Applications of isolated mitochondria have shown to be advantageous in the models of spinal cord, lung, kidney and brain injuries (62). Moreover, it has been reported that platelet-derived mitochondria transfer facilitated wound-closure by modulating ROS levels in dermal fibroblasts (63). Interestingly, Wu et al. (2024) engineered nanomotorized mitochondria with chemotactic targeting capabilities for damaged heart tissue, encapsulated in enteric capsules to protect them from gastric acid erosion prior to oral administration (64). They discovered that once released in the intestine, the mitochondria were quickly absorbed by intestinal cells and entered the bloodstream, indicating their rapid uptake by the intestinal cells and distribution *in vivo*. Additionally, several studies have shown that mitochondria from human MSCs can enter rodent cells *in vitro* [reviewed in (52, 65, 66)] or *in vivo* after injection into rodent tissues (52, 67–71).

While the underlying mechanisms remain to be fully explored *in vivo* in this study, our *in vitro* data—obtained in parallel using the same batches of “freshly” isolated mitochondria used for *in vivo* transplantation—demonstrate a dose-dependent increase in viable MitoTracker Red+ mitochondria, enhanced metabolic activity, and elevated ATP content in colonic recipient cells. These *in vitro* findings support the observed *in vivo* therapeutic effects following local administration. Masuzawa et al. observed that autologous mitochondrial transplantation led to the internalization of mitochondria by cardiomyocytes, resulting in enhanced ATP production and an increase in differentially expressed proteins related to mitochondrial pathways and cellular respiration (72). Recent reviews have comprehensively covered the molecular mechanisms and therapeutic potential of MSCs as a source for mitochondrial transplantation across various diseases (27, 73). Future *in vivo* studies using pre-labeled human mitochondria should enable tracing of their post-transplantation distribution, biogenesis, functionality, and any risks associated with their *in vivo* toxicity in different models of digestive fistula.

## Conclusions

Our investigation into mitochondria-based cell-free nano-biotherapy for post-surgical fistulas following laparoscopic sleeve gastrectomy unveils a promising therapeutic avenue amidst the challenges posed by the obesity pandemic. Leveraging the pivotal role of energy metabolism and mitochondrial function in tissue repair, our study successfully isolates structurally intact and functionally viable mitochondria from hMSCs. Through *in vitro* test, we demonstrate the significant metabolic effects of isolated mitochondria on the starved model of colonic epithelial cells. Our findings particularly underscore the potential of allogenic mitochondria transplantation to mitigate postoperative gastric fistula, offering a novel and promising approach for enhancing patient outcomes in bariatric surgery and beyond.

## Abbreviations

hMSCs: Human mesenchymal stem/stromal cells
HCEC-1CT: Human colonic epithelial cells
EV: Extracellular vesicles
ECMO: Extracorporeal membrane oxygenation
MHC: Major histocompatibility complex
α-MEM: α-Minimal Essential Medium
FBS: Fetal bovine serum
PBS: Phosphate-buffered saline
EDTA: Ethylenediaminetetraacetic acid
SN: Supernatant
Mito: Isolated mitochondria
ILM: Interferometric light microscopy
Part/mL: Particles per milliliter
SEM: Scanning electron microscopy
TEM: Transmission electron microscopy
OXPHOS: Oxidative phosphorylation
POD: Post-operative day(s)
RB: Respiration buffer
LSG: Laparoscopic sleeve gastrectomy
PRP: Platelet-rich plasma

## Declarations

### Ethics approval and consent to participate

All experiments were approved by the animal care and use committee in France and the Ministry of Higher Education and Research (APAFIS #35921-2022010121242601).

### Consent for publication

Not applicable.

### Availability of data and materials

The datasets used and/or analyzed during the current study are available from the corresponding authors on reasonable request.

### Author’s contribution

Conceptualization: A.M., A.K.A.S., G.R., S.M.; formal analysis: A.M., A.G., A.C.S., S.M.; methodology: A.M., A.G., A.C.S., A.C.A., Z.A.A.D, C.R., D.A., S.M.; project administration: A.M, G.L., F.G., A.K.A.S, S.M.; resources: M.K., G.L., F.G., A.K.A.S., G.R., S.M.; supervision: A.K.A.S., G.R., S.M.; validation: A.M., A.G., A.C.S., A.K.A.S., G.R., S.M. writing—original draft: A.M., A.G., A.C.S, S.M.; writing—review and editing: A.M., A.G., A.C.S, A.C.A., Z.A.A.D., M.K., G.L., F.G., A.K.A.S., G.R., S.M. All authors have read and agreed to the published version of the manuscript.

## Acknowledgments

This work was supported by the IdEx Université Paris Cité, ANR-18-IDEX-0001 (IVETh plateform), by the Region Ile de France under the convention SESAME 2019 - IVETh (EX047011) (IVETh plateform) and by the Region Ile de France and Banque pour l’Investissement (BPI) under the convention Accompagnement et transformation des filières projet de recherché et développement N° DOS0154423/00 & DOS0154424/00 (IVETh platform). This project has received funding from CNRS and from the European Research Council (ERC) under the European Union’s Horizon 2020 research and innovation program (grant agreement No.852791). We would like also to acknowledge technical assistance from the Microscopy and Imaging Platform MIMA2 (INRAE, Jouy-en-Josas, France) DOI: MIMA2, INRAE, 2018; Microscopy and Imaging Facility for Microbes, Animals and Foods. In particular, we would like to thank Christine Longin and Vlad Costache from the INRAE MIMA2 platform for their valuable help in performing TEM and SEM analyses. The authors declare that artificial intelligence is not used in this study.

