## Supplementary Figure 1 for "Therapeutic potential of human mesenchymal stromal cell-derived mitochondria in a rat model of post-surgical digestive fistula: towards an energetic nano-biotherapy"

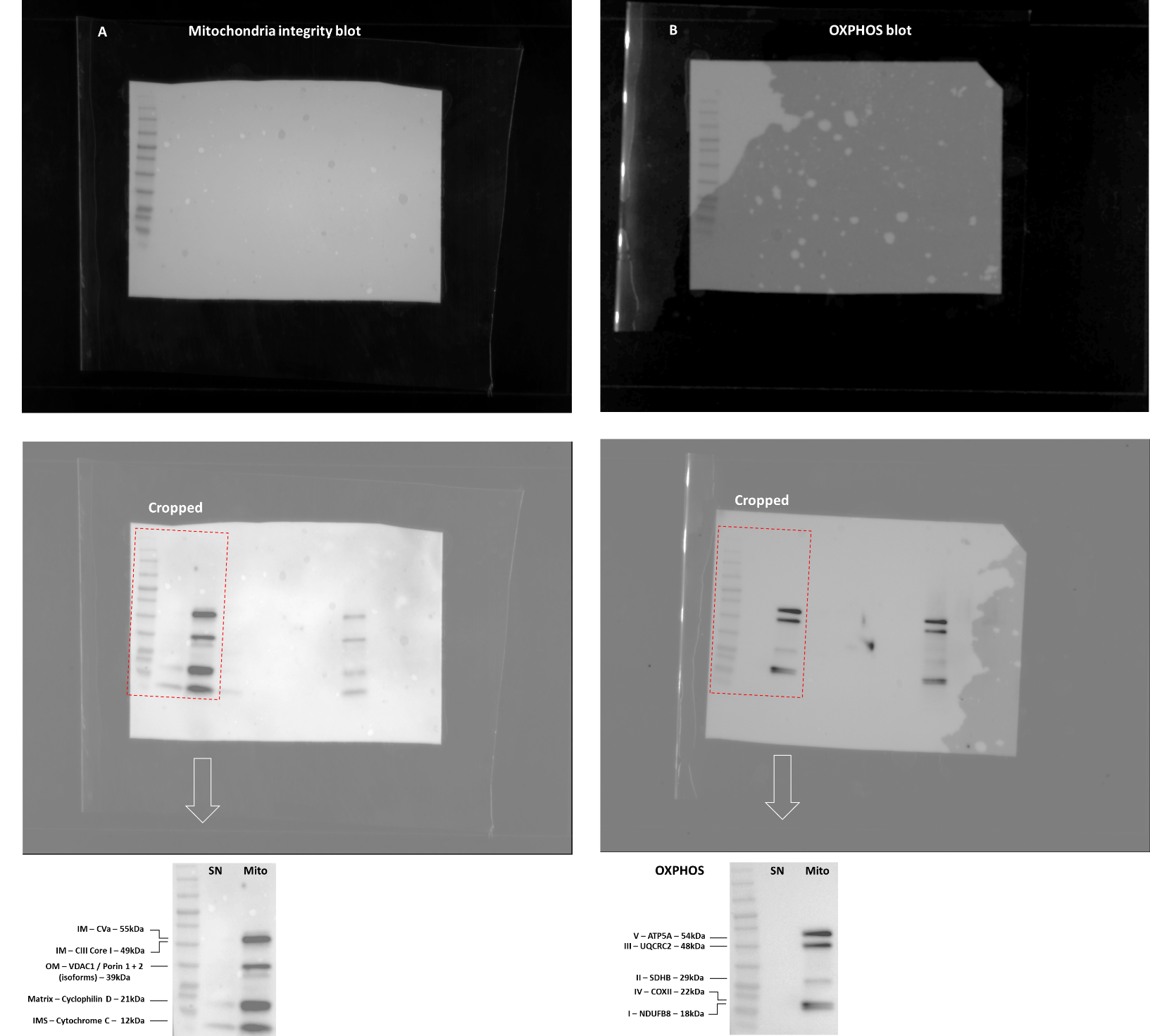
**Supplementary Figure 1**

***Supplementary Figure 1.*** *The uncropped full-length blots. To improve the clarity and conciseness of the presentation the blots of the mitochondria membrane integrity as well as OXPHOS western blotting were cropped. (A and B) shows these uncropped images before and after immunodetection. The cropped blots containing the band sizes and labels are found in the lower panels as well as in the Figure 1F-G.*
